# Phylogenetic expression profiling reveals widespread coordinated evolution of gene expression

**DOI:** 10.1101/045211

**Authors:** Trevor Martin, Hunter B. Fraser

## Abstract

Phylogenetic profiling, which infers functional relationships between genes based on patterns of gene presence/absence across species, has proven to be highly effective. Here we introduce a complementary approach, phylogenetic expression profiling (PEP), which detects gene sets with correlated expression levels across a phylogeny. Applying PEP to RNA-seq data consisting of 657 samples from 309 diverse unicellular eukaryotes, we found several hundred gene sets evolving in a coordinated fashion. These allowed us to predict a role of the Golgi apparatus in Alzheimer’s disease, as well as novel genes related to diabetes pathways. We also detected adaptive evolution of tRNA ligase levels to match genome-wide codon usage. In sum, we found that PEP is an effective method for inferring functional relationships—especially among core cellular components that are never lost, to which phylogenetic profiling cannot be applied—and that many subunits of the most conserved molecular machines are coexpressed across eukaryotes.

## Introduction

Many cellular functions are carried out by groups of proteins that must work together, such as pathways and protein complexes. When one of these functions is no longer needed by a particular species, then there is no longer any selection to maintain the genes needed specifically for this function, and they will eventually deteriorate into pseudogenes or be lost altogether. A method known as phylogenetic profiling (PP) leverages this idea, correlating patterns of gene presence/absence across species to identify functionally related genes (Pellegrini et al. 1999). For example, this technique has been used to discover novel genes involved in Bardet-Biedl Syndrome (BBS) (Mykytyn et al. 2004; Chiang et al. 2004; Li et al. 2004) and mitochondrial disease (Pagliarini et al. 2008), since these diseases involve genes that have been lost in multiple independent lineages. In these studies, patterns of gene conservation across species are typically represented by their binary presence/absence, and knowledge of the species phylogeny is used to identify genes whose losses have coincided with those of well-characterized genes (Li et al. 2014). Coordinated gene losses can then be analyzed for gene pairs individually or gene groups as a whole to reveal functional relationships (Tabach et al. 2013b).

In addition to the correlated gene losses that are the focus of phylogenetic profiling (PP), coordinated evolution of gene expression levels can also indicate functional similarity (in this work we distinguish between coevolution, in which a gene evolves in direct response to changes in another, and coordinated/correlated evolution, in which genes evolve in a coordinated fashion that may or may not be in response to one another). For example, coordinated evolutionary changes have been observed between computationally predicted expression levels (based on codon usage bias) in yeast and other microbes (Lithwick and Margalit 2005; Fraser et al. 2004). Experimentally measured gene expression levels could also potentially uncover genes with correlated evolution, including genes that are never lost and thus not amenable to PP (Figure 1A); however in practice this has not been possible because of the small number of species, and the narrow phylogenetic breadth, in previous studies of gene expression evolution. The largest such studies have been limited to a few dozen species and have focused exclusively on mammals (Perry et al. 2012; Fushan et al. 2015) or yeast (Thompson et al. 2013), in contrast to recent PP studies that rely on hundreds of complete genome sequences from widely divergent species (Li et al. 2014; Dey et al. 2015; Tabach et al. 2013a).

**Figure 1.**
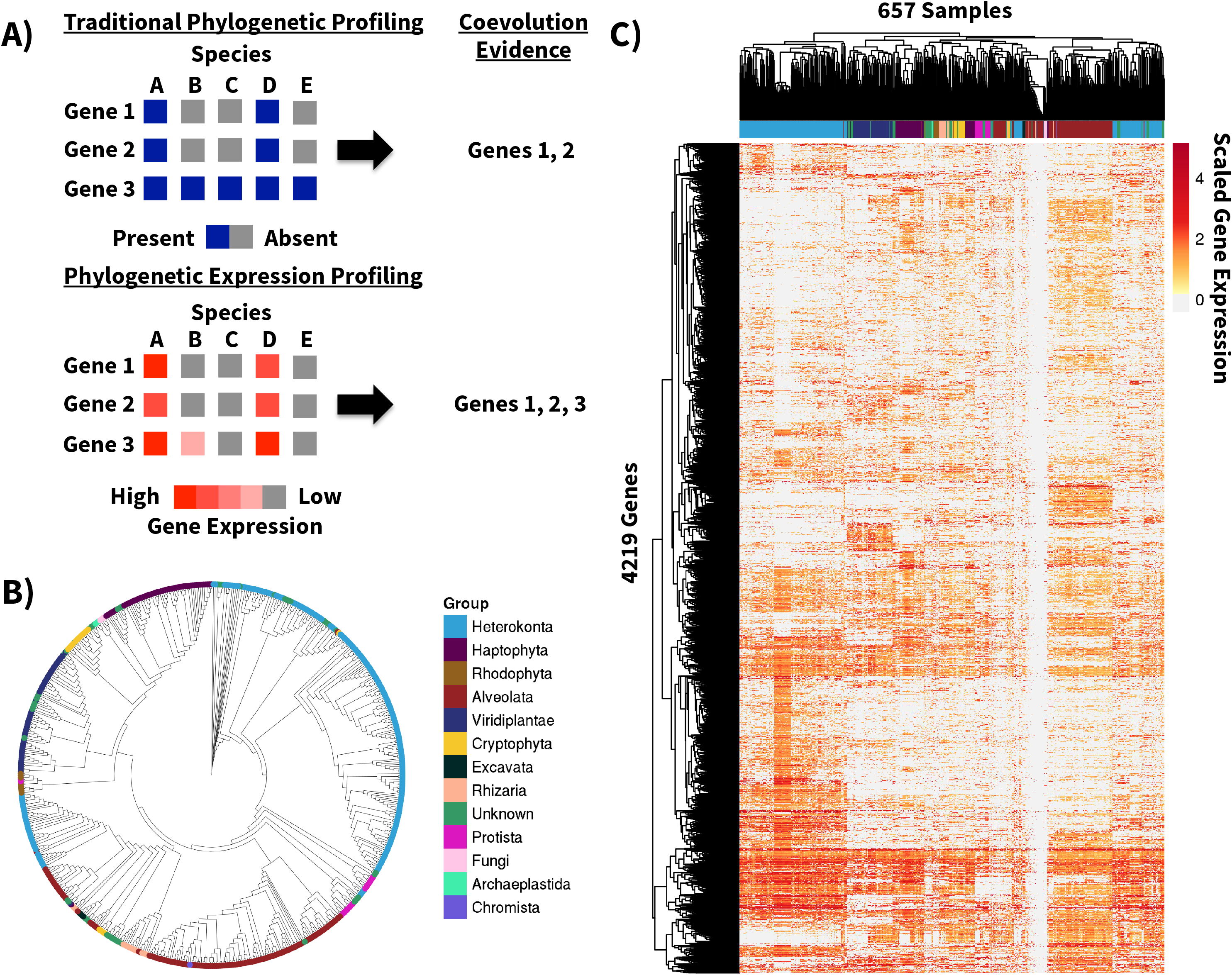
Overview of phylogenetic expression profiling approach and data. **(A)** Overview of traditional phylogenetic profiling (PP; top) and phylogenetic expression profiling (PEP; bottom). PEP uses the quantitative gene expression levels across species rather than the binary presence/absence of a gene. Patterns of coordinated evolution hidden to PP can be potentially uncovered using PEP. **(B)** 18S-based cladogram of the species in this study. **(C)** Heatmap of expression levels of the 4,219 genes and 657 samples analyzed in this study. Samples and genes are clustered hierarchically. Color bar near the top shows the phylum of each sample.

In this study we developed an approach, phylogenetic expression profiling (PEP), that utilizes cross-species gene expression data to infer functional relationships between genes. We applied PEP to the most phylogenetically diverse gene expression data set generated to date: RNA-seq for 657 samples from 309 species. These species are eukaryotic marine microbes collected from across the world, spanning 12 phyla that represent most major eukaryotic lineages, including many rarely studied clades that lack even a single sequenced genome (Figure 1B) (Keeling et al. 2014). All RNA samples were prepared, sequenced, and analyzed following a standardized pipeline established by the Marine Microbial Eukaryotic Transcriptome Project (MMETSP) (Keeling et al. 2014). Some of the MMETSP RNA-seq data has been examined in studies of specific species (Ryan et al. 2014; Koid et al. 2014; Santoferrara et al. 2014; Frischkorn et al. 2014), but the data have not previously been analyzed collectively.

## Results

In order to apply PEP to the MMETSP data (Keeling et al. 2014), we created a matrix of gene expression levels for 4,219 genes that had detectable expression in at least 100 of the 657 samples (Figure 1C; Supplemental Fig. S1; see Methods). To identify coordinated evolution, we calculated all pairwise Spearman correlations between genes in the expression matrix. Because of the complex phylogenetic structure of the data, which can inflate correlations due to non-independence, we did not attempt to assign p-values to individual pairwise PEP correlations; rather we focused on detecting coordinately evolving groups of genes, for which we can create a random permutation-based null distribution that precisely captures the effects of phylogenetic structure, even when the phylogeny is not known (Fraser 2013) (see Methods). To ensure that PEP does not utilize gene presence/absence information—so that its results are independent of PP—we restricted all correlations to samples in which a given pair of genes were both detectably expressed.

To test the performance of PEP, we compared our results to PP in two ways. First, we examined genes with a known role in cilia, since this organelle is one of the most significant gene sets implicated by many PP studies (Dey et al. 2015; Li et al. 2014; Avidor-Reiss et al. 2004). We found this gene set was also enriched for high PEP correlations (Figure 2A; p = 5.1x10^−5^), indicating that the ciliary genes show coordinated evolution of gene expression, in addition to gene loss. We then compared the two methods at a finer scale, by asking whether the specific ciliary gene pairs with the strongest PP signal also show coordinated evolution by PEP. Comparing the PEP correlations to the binary presence/absence PP correlations, we found a moderate level of agreement (Figure 2B; p = 3.4x10^−22^), suggesting that the specific ciliary gene pairs most likely to be lost together also tend to have coordinately evolving expression levels.

**Figure 2.**
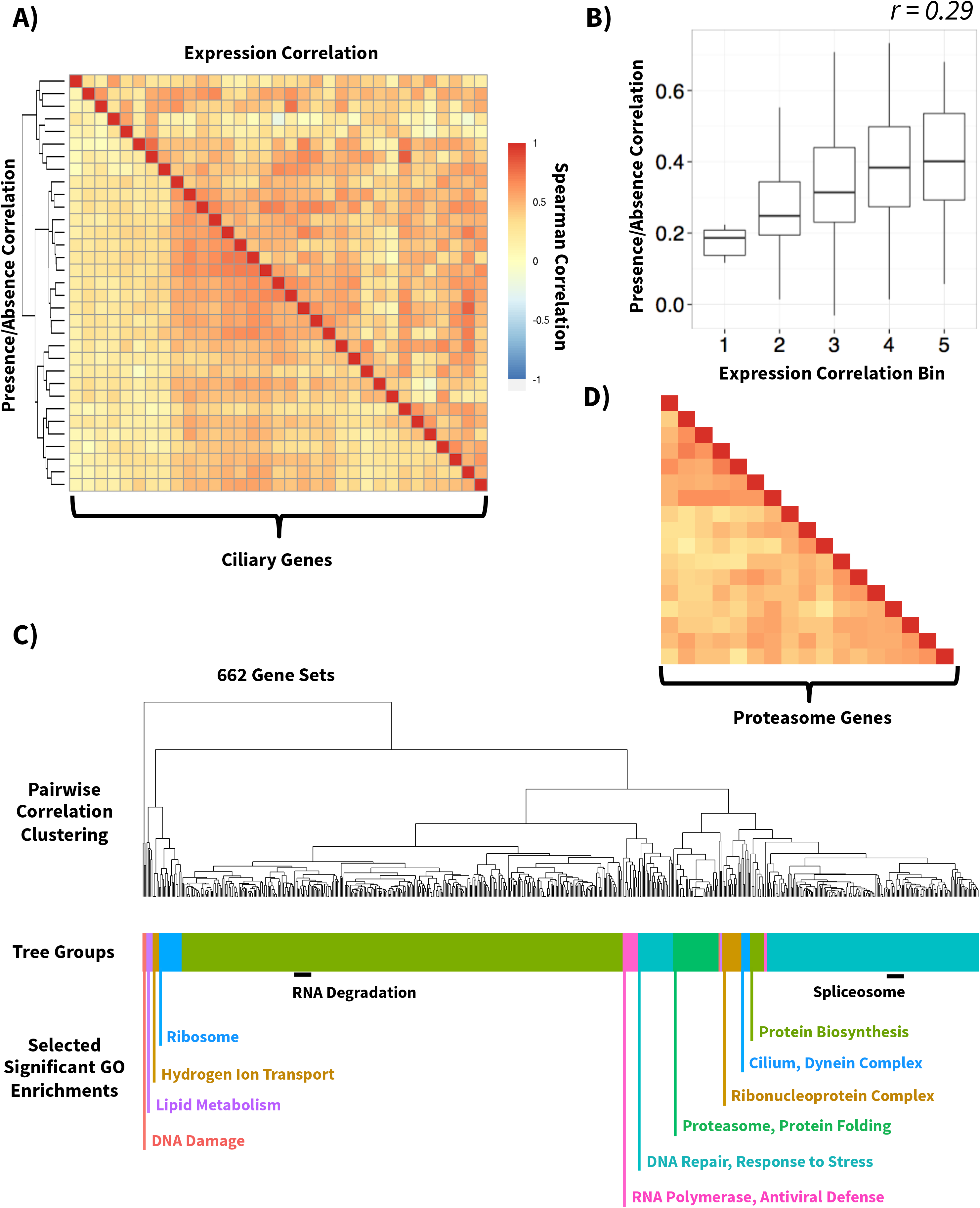
Phylogenetic expression profiling reveals coordinated evolution within gene sets. **(A)** Heatmap of the PP (presence/absence, bottom left) and PEP (expression, top right) Spearman correlation based scores (see Methods) between ciliary genes. Both PP and PEP values are hierarchically clustered by the Euclidean distance between the gene PP scores. **(B)** For ciliary genes in part (A), pairwise PP correlations increase with PEP correlation strength. **(C)** The 662 gene sets with significant PEP scores are clustered by the pairwise correlations between gene sets. The color bar below the dendrogram shows the 15 unique gene set groups the dendrogram was divided into and gene ontology enrichments for each group are highlighted in the same color. Black bars highlight notable groups of gene sets within the larger groups. **(D)** Proteasome genes were found to be undergoing coordinated evolution and are shown as a heatmap with the same scale for the gene-gene scores as in (A).

We then asked whether PEP and PP agree at a more broad scale, by testing whether the collection of all coordinately evolving modules identified in a recent PP study (Li et al. 2014) showed increased PEP signal as well. To achieve this, we calculated the median PEP score within each of the 327 PP modules. We found significant (p = 2.9x10^−47^; see Methods) enrichment for PEP correlations in these previously identified PP modules, suggesting that PEP detects many of the same gene sets implicated by PP. For example, some of the strongest PEP correlations were for gene sets involved in the ribosome, spliceosome, and cilia.

To identify additional coordinately evolving modules not detected by PP, we applied PEP to a collection of 5,914 previously characterized gene sets, including both pathway and disease databases (see Methods). Of these, we found 662 gene sets with significant coordinated evolution, compared to only ~33 expected at this level by chance (Figure 2C; 5% FDR; Supplemental Table S1). Most of these were gene sets with no previous evidence of coordinated evolution from PP studies, such as RNA degradation, the proteasome, and the nuclear pore complex. Examining all pairwise PEP correlations within each of these gene sets revealed that the coordinated evolution tends to be shared across most gene pairs, rather than only driven by a small subset of them (e.g. as shown for proteasome genes in Figure 2D). Many of these coordinately evolving gene sets have been hidden from PP analysis because of the rarity of losing these genes (Supplemental Fig. S2).

Having identified hundreds of cases of coordinated evolution within gene sets, we then asked whether we could also detect coordinated evolution between gene sets. To identify these, we calculated the PEP correlation between each pair of genes in a given pair of gene sets (excluding any genes present in both; see Methods). Among the 218,791 pairs of gene sets we tested, 22,665 had evidence of coordinated evolution (with <1 expected by chance; Supplemental Table S2). For example, we found that genes involved in the Golgi apparatus had strong evidence (p = 2.9x10^−5^) of coordinated evolution with genes down-regulated in Alzheimer’s disease (Figure 3A). Previous studies have implicated Golgi fragmentation in the pathogenesis of Alzheimer’s (Joshi et al. 2014; Joshi et al. 2015) and this coordinated evolution gives additional evidence for a functional relationship between these gene sets.

**Figure 3.**
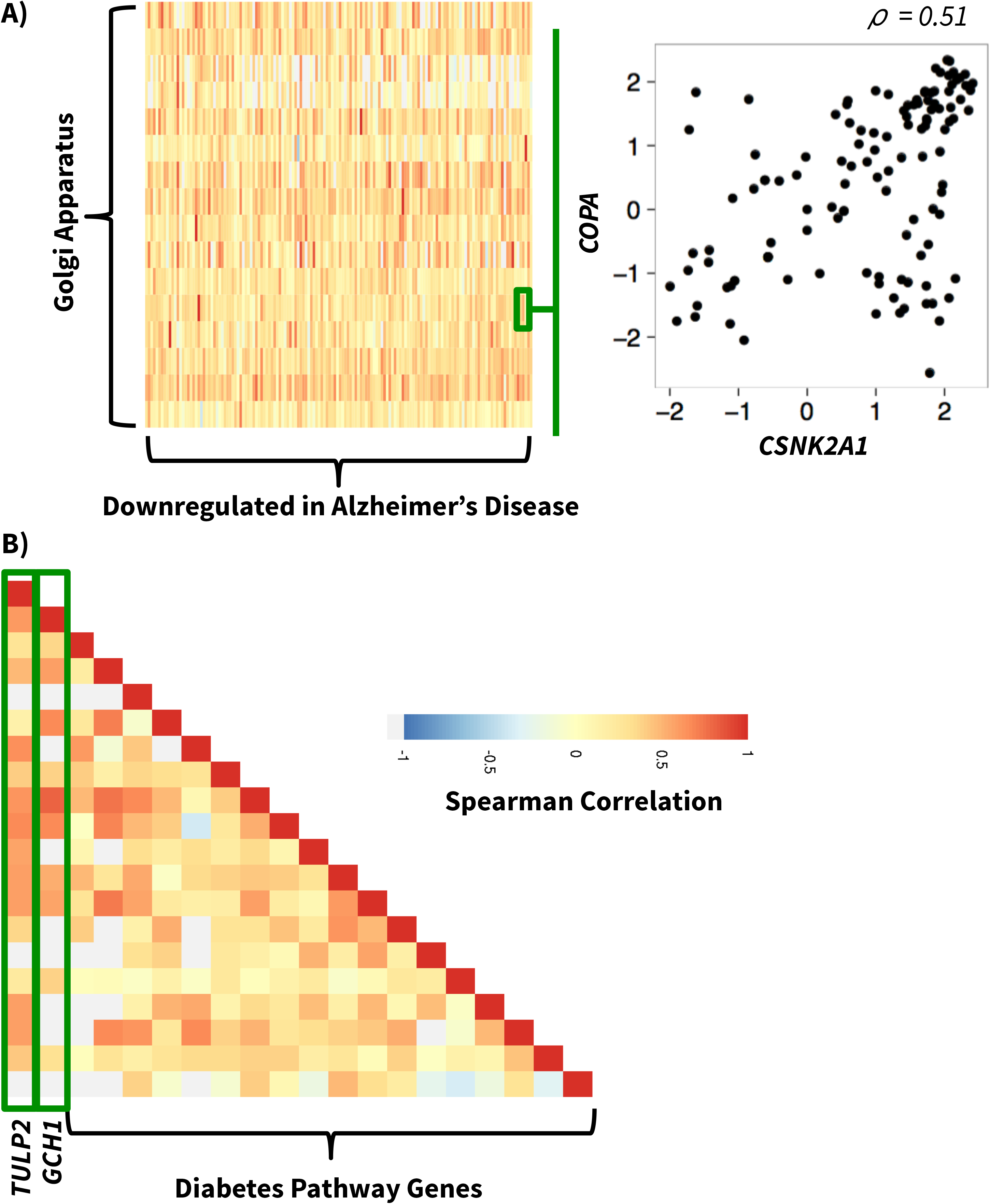
Coordinated evolution between gene sets and addition of novel genes to known gene sets. **(A)** The coordinated evolution scores between the gene sets for the Golgi apparatus and genes downregulated in Alzheimer’s disease is shown as a heatmap with the same scale as in (B). The gene pair highlighted in green is shown as a scatterplot to the right; each point is a sample with measured expression. **(B)** The coordinated evolution scores for diabetes pathway genes are shown in heatmap form. In green are the two genes not in this gene set with the strongest PEP scores to the known genes in this set.

In addition to identifying coordinated evolution within and between known gene sets, PEP can also implicate novel genes evolving in tandem with a known gene set. For this analysis, we calculated the PEP correlation between the genes in a given set and every other gene; those with the strongest median correlations are most likely to be functionally related to that set. Of particular interest is identifying novel disease-related genes by implicating genes related to disease pathways. For example, *TULP2*—a member of the tubby-like gene family—had the highest PEP correlation with the diabetes pathway gene set (Figure 3B; p = 3.0x10^−4^) and is in a linkage region for severe obesity (Bell et al. 2004). The second strongest PEP correlation (p = 9.1x10^−3^) is *GCH1*, which is adjacent to a SNP with moderate (p = 6.1x10^−6^) association with type 2 diabetes (T2D) (Wellcome Trust Case Control Consortium 2007). Moreover, GCH1 contains SNPs strongly (p = 7.6x10^−64^) associated with circulating galectin-3 levels, which is itself associated with insulin resistance and obesity (de Boer et al. 2012). The genes coordinately evolving with diabetes pathways were enriched for T2D GWAS associations (p = 2.3x10^−2^ for the top 10 genes, and p = 2.6x10^−2^ for the genome-wide trend; see Methods), suggesting that the PEP correlations are indeed predictive of genes involved in T2D.

Several characteristics of the MMETSP samples, such as the location of collection, were recorded for most samples. To investigate whether any gene expression levels show a latitudinal gradient across the diverse set of MMETSP species, we correlated absolute latitude with expression levels of every gene. Although we did not find any functions significantly enriched in the latitude-associated genes, the most strongly correlated gene was the translesion DNA polymerase *POLH* (Figure 4A; Spearman’s ρ = -0.52, p = 2.4x10^−24^). Expression was generally higher closer to the equator, as expected if its mRNA level has evolved in response to the local levels of UV radiation.

**Figure 4.**
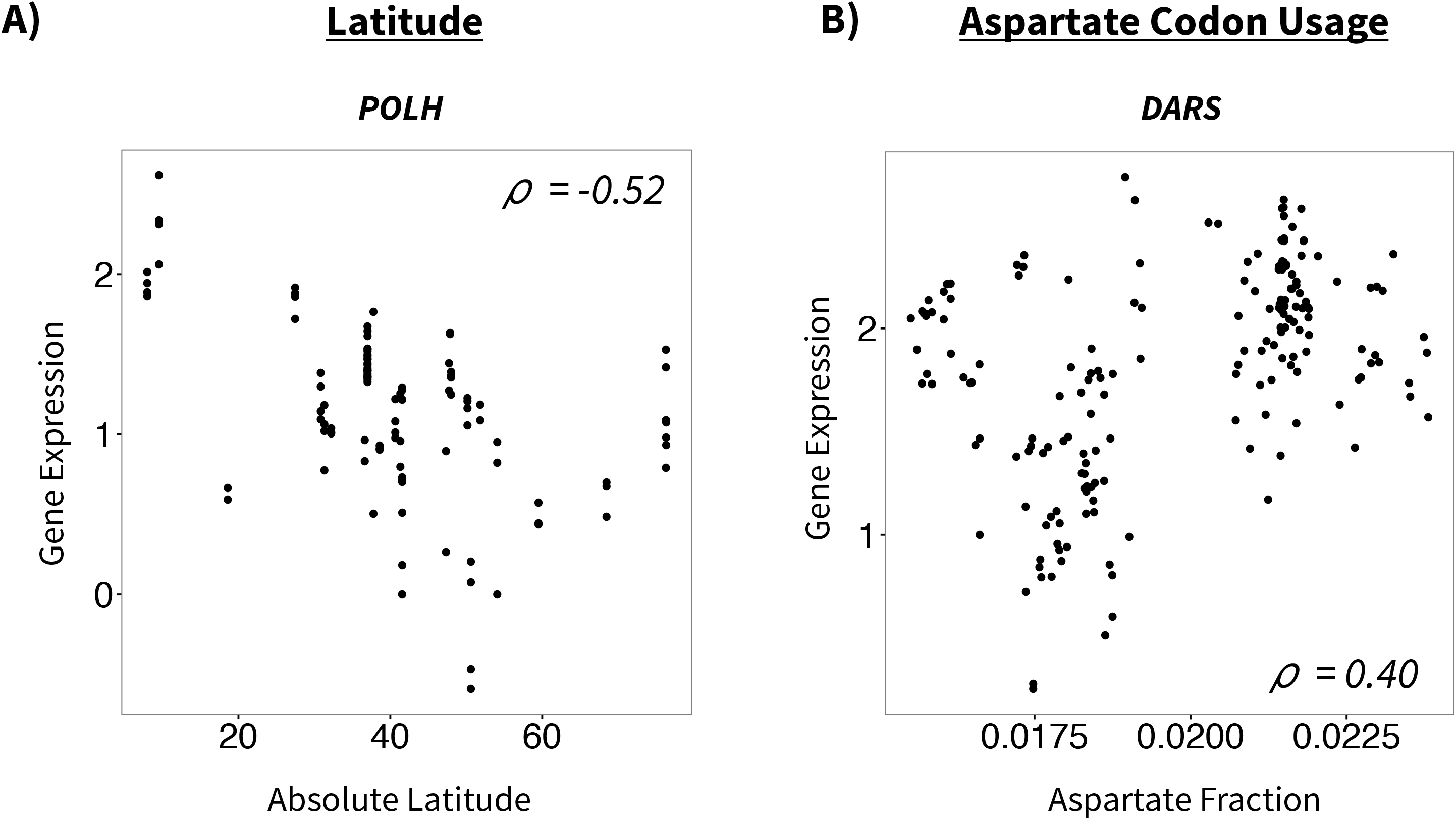
Associations between expression and other sample information. **(A)** Scatterplot of the association between gene expression and the absolute value of latitude for the DNA polymerase *POLH*. Each point represents a sample with measured expression. **(B)** Scatterplot of the association between expression of the aspartate-tRNA ligase *DARS* and the abundance of aspartate codons in the coding regions of each sample’s transcriptome.

Other characteristics of each species can be estimated directly from the assembled transcriptomes. For example, in each species we calculated the genome-wide fraction of codons encoding each amino acid, and tested whether these fractions predict the expression levels of the corresponding tRNA ligases—enzymes that “charge tRNAs with the appropriate amino acid”. Of the ten tRNA ligases with expression data, all ten had a higher than codon median correlation with the relative abundances of their respective codons (binomial p = 9.8x10^−4^). For example, the association between the expression of the aspartate-tRNA ligase (DARS) and aspartate codon abundance is shown in Figure 4B (Spearman’s ρ = 0.40, p = 4.8x10^−15^).

## Discussion

The PEP method introduced here builds on traditional PP, and together with the most phylogenetically diverse gene expression data set available to date, it revealed widespread evidence for coordinated evolution of gene expression. Interestingly, our method – which only relies on species where a gene is present – identified previously well-studied gene sets with coordinated losses such as cilial genes, in addition to identifying many previously unidentified sets of coordinatedly evolving genes. One explanation for why gene sets such as mismatch repair have not been identified by PP is that PP is substantially underpowered to detect these gene sets because of the rarity of their loss across species.

Further, our analysis of coordinated evolution between gene sets allows us to infer functional linkages between known biological pathways. In particular, Golgi fragmentation in Alzheimer’s disease has been linked to promotion of amyloid beta production (Joshi et al. 2014) and potential phosphorylation of the tau protein which underlies the formation of neurofibrillary tangles (Jiang et al. 2014). Notably, since the Golgi genes are not themselves mis-regulated in Alzheimer’s disease, a differential expression analysis of the Alzheimer’s patient samples vs. controls could not provide this connection. Additionally, genes such as *GCH1* with nominally but not genome-wide significant p-values of association with traits such as T2D would not be identified as playing a role in disease pathways without the orthogonal evidence of association such as the coordinated evolution evidence presented in this study.

Previously, within single species or genera, many latitudinal gradients of traits have been reported, which are often attributed to local adaptations to climate (Fraser 2013; Hancock et al. 2011; Franks and Hoffmann 2012; Savolainen et al. 2013). This study has expanded such analyses across major phylogenetic groups and the identification of *POLH*, which plays an important role in the repair of UV-induced damage and leads to xeroderma pigmentosum when mutated in humans (Masutani et al. 1999), is a novel example of a potential gene expression based response to environmental variability. Additionally, our identification of tRNA ligase levels as associating with codon abundance suggests that tRNA ligase levels may adaptively evolve in response to species-specific codon usage, and is consistent with patterns of tRNA gene copy number and codon usage in bacteria (Higgs and Ran 2008).

Overall, applying our PEP framework to a gene expression data set of unprecedented phylogenetic diversity, we identified many novel examples of coordinated evolution. These included hundreds of cases of coordinated evolution within and between previously characterized gene sets, and also coordinated evolution implicating novel genes related to diabetes pathways. Although it may at first seem surprising that unicellular eukaryotes could shed light on complex diseases like Alzheimer’s and T2D, the fact that the genes are present throughout eukaryotes suggests that the underlying cellular functions are far more conserved than the specific human disease phenotypes—consistent with previous work, for example using yeast to study Parkinson’s disease (Gitler et al. 2008) and plants to study neural crest defects (McGary et al. 2010).

We expect that as the diversity of species with publicly available gene expression data continues to grow, PEP will become a powerful approach for detecting coordinated evolution at the molecular level, and for leveraging these patterns to inform us about functional connections between genes conserved throughout the tree of life.

## Methods

### Data filtering and normalization

Raw reads and transcriptome assembled coding sequence (CDS) data for 669 individually annotated samples and 119 jointly annotated sample sets from the Marine Microbial Eukaryote Transcriptome Sequencing Project (MMETSP) were downloaded from the CAMERA database (http://camera.crbs.ucsd.edu/mmetsp/index.php). Details on each annotation method (performed by the MMETSP project) can be found on the MMETSP website (http://marinemicroeukaryotes.org/resources). All raw reads were then normalized using the Transcripts per Million (TPM) normalization technique (Wagner et al. 2012). Up to five Swissprot ID annotations provided by MMETSP for each CDS were then culled for IDs that had a BLASTP alignment score of at least 80% the maximum alignment score for that CDS, and converted to UniRef100 IDs (UniRef IDs are comprehensive non-redundant clusters of UniProt sequences (Suzek et al. 2007)). In order to create vectors of expression across samples for a set of UniRef100 defined “genes”, normalized read counts were combined within a sample for CDSs that had at least three matching UniRef100 IDs and then across samples by ranking the UniRef100 IDs by alignment score and then creating an expression vector for an annotation by matching the unique top ranked annotations against the top ranked annotation for each CDS in each sample with ties resolved by annotation score. This initial round of matching was then followed by matching successively lower ranked annotations of still unmatched normalized read counts until all are combined into expression vectors. For details see Supplemental Fig. S1. The expression vectors with at least 100 samples with measured expression were then combined into a matrix with each column as a sample (669 and 387 samples, for the individual and jointly annotated sets respectively) and each row as a gene (4,995 and 1,051 genes).

For the individually annotated samples, this expression matrix was then normalized by dividing each sample by the total number of genes in that sample and then adjusting for batch effects by regressing out the MMETSP transcriptome pipeline used (the two pipelines used differed in the method for transcriptome assembly), the day the sample was processed, and the lab that submitted the sample, setting the value of samples missing expression to zero for the regression step. All variables were regressed out as binary factors. Any samples missing any of these variables were dropped from the analysis.

For both sample annotations, the UniRef100 ID for each gene was converted to a UniRef50 ID (a more lenient across-species gene clustering than the UniRef100 ID) and any expression vectors with the same ID were collapsed by sum. The resulting individual annotation matrix has 4,219 genes and 657 samples and the combined annotation matrix has 1,031 genes and 387 samples.

### Phylogenetic expression profiling

Gene sets from the Online Mendelian Inheritance in Man (OMIM) database (McKusick-Nathans Institute of Genetic Medicine, Johns Hopkins University (Baltimore, MD)), the Human Phenotype Ontology (HPO) database (Kohler et al. 2014), the Mouse Genome Informatics (MGI) database (Eppig et al. 2015), and the Molecular Signatures Database (MSigDB) (Subramanian et al. 2005) were downloaded to create a list of 5,914 gene sets with at least three genes that mapped to UniRef50 IDs in the individual annotation data set.

Phylogenetic expression profiling (PEP) tests for coordinated evolution of gene expression levels by calculating the median Spearman correlation between all pairwise combinations of genes in a gene set. Importantly, each pairwise correlation was calculated using only the samples that had expression measured for each gene and the genes that had at least 20 such samples. To calculate the significance of this median correlation, it was compared to 10,000 null median correlations created by random gene sets with the same number of genes, drawn from the 25 genes that most closely match the data missingness profile of the gene they replace. The data missingness profile for a gene pair was quantified by the Euclidean distance between the presence/absence vector of each gene across samples. The significance was then given by:

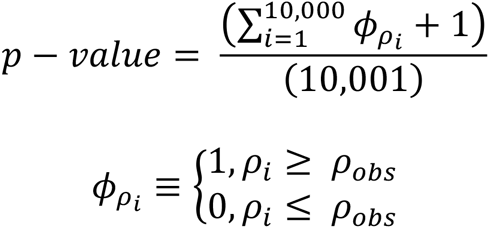

The false discovery rate (FDR) is then determined by treating each of the 10,000 permutations as the real data and calculating 10,000 sets of p-values as above. A sliding p-value cutoff is then instituted and the ratio of p-values below this cutoff in the real data to the mean of the number of p-values below this cutoff in the 10,000 null permutations is the FDR.

### Comparison with previous methods

Evolutionarily conserved modules (ECMs) from the clustering by inferred models of evolution (CLIME) algorithm applied to human pathways were downloaded from the CLIME website (http://www.gene6clime.org/). The 327 ECMs with an ECM score of greater than five and at least two genes in the individual annotation matrix were used for the validation test. To validate the PEP method, we calculated the median correlations for these ECMs in the same way as PEP, and the median of this distribution across ECMs was then compared to 10,000 null medians calculated using the same null strategy as PEP. The permuation p-value for enrichment for high PEP scores is then:

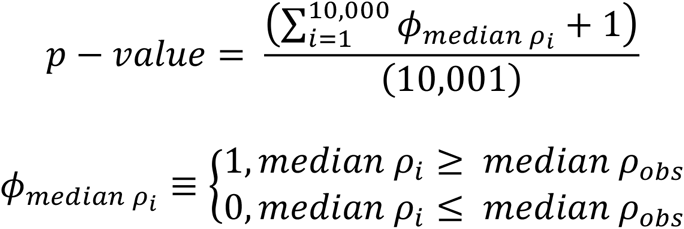

Since the observed statistic was more extreme than all 10,000 permutations, a z-score based p-value was estimated:

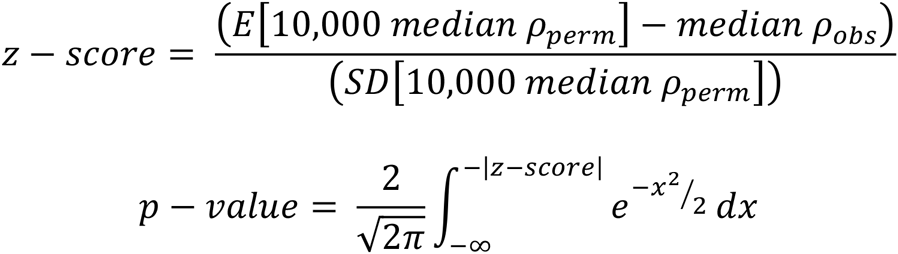

The PP correlation of a gene set was calculated by taking the median of the Pearson correlation of each pairwise presence/absence vector for each gene in the set. For a gene set, the PP correlation was then compared to the PEP correlation for each gene by calculating the Pearson correlation between the PEP and PP correlations for each gene pair. The significance of this correlation was then calculated by permuting the presence and absence vectors for each gene in the set and then recalculating the PEP vs. PP correlation 10,000 times; the number of times a permutation beat or matched the observed value divided by the number of permutations was then the permutation p-value which was then converted to a z-score based p-value as above.

### Phylogenetic tree construction

The 18S sequence available for 655 samples was downloaded from the CAMERA database as above and aligned using the multiple sequence alignment tool Clustal Omega (Sievers et al. 2011). This alignment was then used to create a maximum likelihood based tree using the program RaxML (Stamatakis 2006) with parameters: -f a –x 12345 –p 12345 -# 100 – m GTRGAMMA. 18S sequences that did not have available sample meta data were then dropped, leaving a total of 635 samples.

### Gene set pairwise comparison

The correlation score between two gene sets was calculated by taking the median of the pairwise gene PEP correlations, excluding any genes present in both gene sets. A dendrogram relating the gene sets with significant PEP scores at a 5% empirical FDR as calculated above was created by calculating the matrix of correlation scores between all the significant gene sets, taking the Euclidean distance between the rows of this matrix, and then hierarchically clustering these distances using the complete linkage algorithm in R’s hclust function (Murtagh). Significance of individual gene set pairwise comparisons was calculated in two steps by first computing the p-value for the observed correlation between each of the gene sets in the comparison and 10,000 gene sets matched by phylogenetic profile and size as in the PEP method above. The maximum of these two p-values for random gene set associations was then taken to give the p-value for the gene set comparison.

Subsets of this dendrogram were then created by cutting the tree at the height which gives 15 unique groups. We tested these subsets for gene set enrichments with the DAVID online enrichment tool (Huang da et al. 2009) using all genes in the individual annotation matrix as background.

### Gene expression/environment comparison

Sample meta data was downloaded from the CAMERA database as described above and included data on 12 measured variables (Latitude, Longitude, pH, Temperature, Salinity, etc.). Additionally, using the downloaded CDS data for each sample, we calculated the genome-wide usage of codons encoding each amino acid.

Significance of expression/environment associations was calculated using the combined annotation matrix data and calculating the Spearman correlation between all the samples with both expression and environmental data. These correlations were converted to p-values by permuting the environmental data 10,000 times and calculating the number of permuted correlations with an absolute value greater than or equal to the observed correlation, divided by the number of permutations as described above. These permutation p-values that beat all permutations were then converted to z-score p-values as described above.

### Addition of genes to gene sets

The correlation of a gene with a gene set was calculated by finding the median PEP correlation of the gene with all the genes in the gene set. The significance of this correlation was calculated by finding the median PEP correlation of the gene with 10,000 permuted gene sets, created as described above, and summing the number of permuted medians with a greater correlation and dividing by the total number of permutations.

To look for genome wide association study (GWAS) hit enrichment, a list of GWAS SNPs with p-value less than 0.05 was downloaded from the Genome-Wide Repository of Associations Between SNPs and Phenotypes (GRASP) database (Leslie et al. 2014). This database was then culled for SNPs with a type II diabetes association and GWAS SNPs in genes were matched to genes in this data set using human gene IDs. Enrichment of GWAS hits in the list of genes added to a gene set was calculated by taking the top ten genes by PEP p-value (with secondary ordering by correlation) with the set and comparing these GWAS p-values to 10,000 random samplings of the same number of GWAS p-values, asking how often a set of p-values smaller than all of the observed p-values are found by chance. To look for a genome-wide trend, the list of genes added to a gene set was divided into 1,000 gene bins ordered by the p-value of PEP association and correlation to calculate the percent of human genes in each bin with a GWAS p-value in the database. The absolute value of the Pearson correlation between gene bin and percent GWAS gene was then compared to 10,000 random permutations of gene ordering.

## Data Access

All MMETSP data are available online at http://camera.crbs.ucsd.edu/mmetsp/index.php.

## Acknowledgements

We would like to thank the members of the Fraser Lab and D. Petrov for helpful discussions and advice, and S. Guida for assistance with the MMETSP data.

## Figures

**Supplemental Figure S1.**
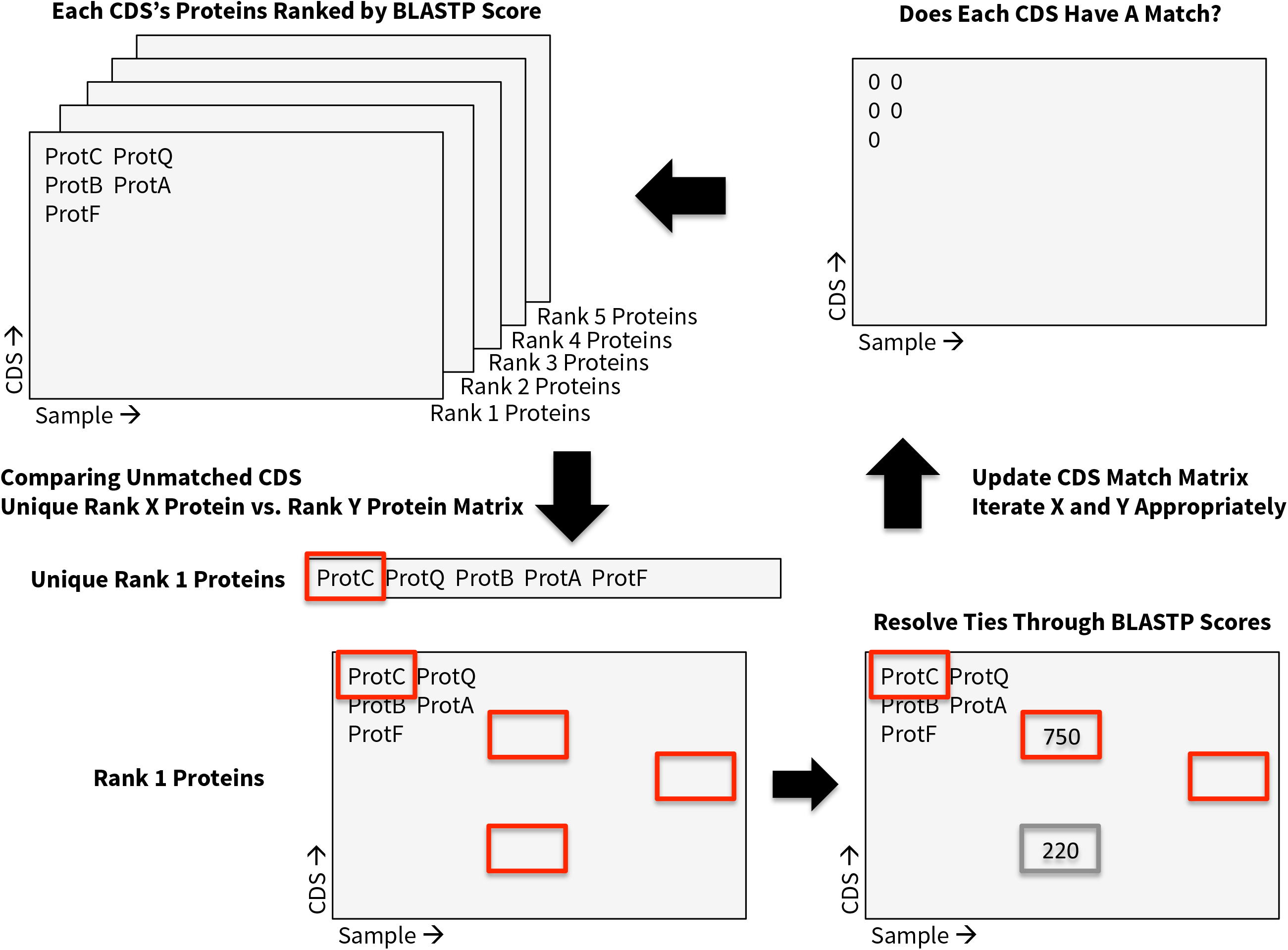
Expression measurements are combined into genes across species based on BLASTP score matching. Each sample has a set of coding sequences (CDS) with measured expression and up to five UniRef IDs identified and ranked by BLASTP score. To begin, all the unique rank one UniRef IDs are used to create a vector of expression for a gene by matching each rank one UniRef ID to the samples with that ID that are also rank one. These samples are then flagged so that expression values are not reused. This process is then repeated by iterating through each set of ranked proteins until all expression values are matched.

**Supplemental Figure S2.**
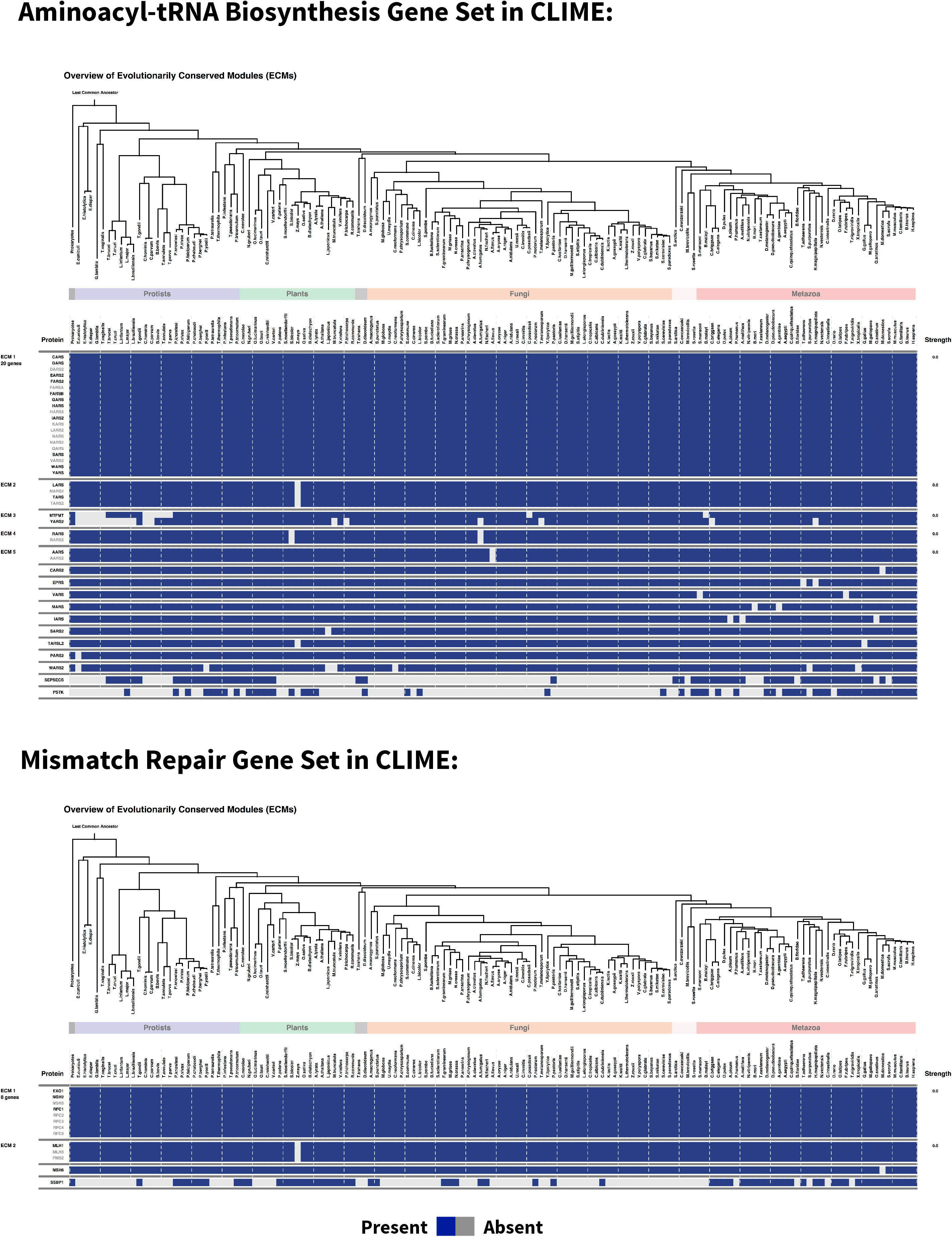
Genes unlikely to be lost are hidden from phylogenetic profiling. Two of the gene sets identified in this study, aminoacyl-tRNA biosynthesis and mismatch repair were also analyzed in a recent PP study (Li et al. 2014) and their patterns of loss over evolutionary time are shown here (blue is gene sequence presence and gray is absence). Neither of these gene sets results in informative signs of correlated loss.

**Supplemental Table S1 Gene sets identified as having significant coordinated evolution.** The name of each gene set with coordinated evolution at a 5% FDR is shown along with the number of genes in that gene set that were present in this analysis, the database source of the gene set, and the nominal p-value of the gene set.

**Supplemental Table S2: Gene set pairs with significant coordinated evolution**.

The name and database source of each gene set pair with coordinated evolution is shown (<1 expected by chance).

